# Integrated Regulation of Dopaminergic and Epigenetic Effectors of Neuroprotection in Parkinson’s Disease Models

**DOI:** 10.1101/2022.06.21.497090

**Authors:** J. Brucker Nourse, Shannon N. Russell, Nathan A. Moniz, Madison Scott, Han-A Park, Kim A. Caldwell, Guy A. Caldwell

## Abstract

Whole exome sequencing of Parkinson’s disease (PD) patient DNA identified single-nucleotide polymorphisms (SNPs) in the *TNK2* gene. Although *TNK2* encodes a non-receptor tyrosine kinase that has been shown to prevent the endocytosis of the dopamine reuptake transporter (DAT), a causal role for TNK2 in PD remains unresolved. We postulated that specific recessive mutations in patients resulted in aberrant or prolonged overactivity as a consequence of failed negative regulation by an E3 ubiquitin ligase, NEDD4. Interestingly, the sole *Caenorhabditis elegans* ortholog of TNK2, termed SID-3, is an established mediator of epigenetic gene silencing and systemic RNA interference facilitated by the SID-1 dsRNA transporter. While SID-3 had no prior association to dopamine neurotransmission in *C. elegans*, we hypothesized that TNK2/SID-3 represented a node of integrated dopaminergic and epigenetic signaling essential to neuronal homeostasis. Using genetic and chemical modifiers, including a TNK2 inhibitor (AIM-100) and NEDD4 activator (NAB2), in bioassays for dopamine uptake or RNAi in dopaminergic neurons of *C. elegans*, we determined that *sid-3* mutants displayed neuroprotection from 6-hydroxydopamine (6-OHDA) exposure, as did wildtype animals treated with AIM-100 or NAB2. Additionally, NAB2 treatment of rat primary neurons correlated with a reduction of TNK2 levels and the attenuation of 6-OHDA neurotoxicity. Notably, CRISPR-modified nematodes engineered with genomic mutations in *sid-3* analogous to PD patient-associated SNPs in *TNK2* circumvented the resistance to RNAi characteristic of SID-3 dysfunction and furthermore exhibited enhanced susceptibility to neurodegeneration. This study describes a molecular etiology for PD whereby dysfunctional cellular dynamics, dopaminergic, and epigenetic signaling intersect to cause neurodegeneration.

**Significance Statement:** The progressive loss of dopamine neurons is a pathological hallmark of Parkinson’s disease (PD). Distinctions between resilience or susceptibility to neurodegeneration in PD are a combined consequence of genetic predisposition and environmental factors, the latter often manifesting as changes in gene expression that are coordinately controlled by small RNA molecules. This research reveals a functional convergence of proteins that modulate uptake of both dopamine and small RNAs, as a regulatory intersection for the integrated control of dopamine neuron health. Analysis of PD-patient mutations in the central protein associated with this functional interface further illustrated the clinical significance of this regulatory mechanism, as well as its potential for therapeutic intervention to prevent neurodegeneration through the fine-tuning of dopamine levels.

## Introduction

Parkinson’s disease (PD) is a neurodegenerative disease associated with the progressive degeneration of dopaminergic neurons in the *substantia nigra*. The significant loss of these neurons results in an attenuated dopaminergic signal, ultimately manifesting into the characteristic symptoms of PD including rigidity and resting tremors. While advances in genome sequencing have expanded the lists of genes and cellular pathways associated with PD, the search to identify a successful disease-modifying therapeutic target remains to be fulfilled. Among these expanding genomic datasets was a whole-exome sequencing (WES) study that revealed the *TNK2* gene, encoding tyrosine non-receptor kinase-2 (*TNK2)*, to be mutated in a group of patients with familial PD (1).

Human *TNK2* (also known as *ACK1*) is involved in several cellular pathways and is highly expressed in pre-synaptic vesicles of neurons in the mammalian brain (2,3), where it functions in the regulation of clathrin-dependent endocytosis of the dopamine transporter (DAT) (4). While the precise mechanism by which TNK2 modulates clathrin-dependent endocytosis is unknown, TNK2 possesses a clathrin-heavy chain domain and has been shown to interact with established regulators of vesicular dynamics including clathrin, adaptor protein-2 (AP-2), and sortin nexin-9 (SNX9) (5). TNK2 is activated by GTP-bound CDC42 phosphorylation, and conversely, is marked for protein degradation by the E3 ubiquitin ligase NEDD4 (neuronal precursor cell-expressed developmentally down-regulated-4) (6). Furthermore, TNK2 specifically interacts with DAT at the neuronal plasma membrane, directly modulating dopamine reuptake from the synapse (7). Our interest in TNK2 was piqued by having previously reported the discovery of a small molecule activator of NEDD4 ubiquitin ligase activity, N-aryl benzimidazole 2 (NAB2), that exhibited potent and selective neuroprotection from dopaminergic neurodegeneration across multiple PD models including in the nematode *Caenorhabditis elegans* and in rat primary neuron cultures, as well as in patient-derived neurons from induced pluripotent stem cells (8,9).

NEDD4 functions to modulate synaptic plasticity in a variety of ways (10), including through the cytoskeletal control of myelin compaction in oligodendrocytes (11), the ubiquitination of AMPARs response to amyloid-beta (Aβ)-peptide induced synaptic dysfunction (12), and in chronic stress response through ubiquitin-targeted degradation of the GluR1 receptor (13). Moreover, NEDD4 appears to be both a target and effector of epigenetic regulation in the chronic stress response, where glutamatergic signaling, DNA methylation, histone modifications, and miRNAs are all suspected determinants of resilience (14). Significantly, *NEDD4* was just recently found to be one of only five genes clinically associated with increased risk of epigenetic modulation in PD (15).

In the aforementioned WES study, four single nucleotide polymorphisms (SNPs) in *TNK2* were identified in PD patients (1). Based on the locations of these SNPs, we hypothesized that PD-associated TNK2 cannot be inactivated or evades being targeted for degradation by NEDD4 ubiquitination, thereby resulting in an improper regulation of DAT, ultimately leading to increased dopamine reuptake and disruption of dopaminergic signaling. Aberrant influx of dopamine not only weakens the synaptic dopaminergic signal but can also be toxic to presynaptic neurons due to dopamine-induced apoptosis (16). We demonstrated a selective vulnerability to dopamine-associated neurotoxicity *in vivo*, using both transgenic worm and mouse alpha-synuclein models of PD, where excess presynaptic dopamine biosynthesis correlated with increased neurodegeneration (17).

The *C. elegans* genome contains a single ortholog to human *TNK2* that is encoded by the *sid-3* (<u>s</u>ystemic RNA<u>i d</u>efective-<u>3</u>) gene. The SID-3 protein is involved in modulating epigenetic signaling through regulating the cellular entry of double-stranded RNA molecules (dsRNAs), such as microRNAs (miRNAs) (18). In worms and mice, respectively, SID-3 and TNK2 have been shown to be required for certain viral-RNA infections as demonstrated by genetic knockout studies (19; 20). The cells of *C. elegans* import dsRNA through the transmembrane transporter SID-1 (21), which is regulated by clathrin-dependent endocytosis (22). Whereas the understanding of organismal dsRNA transport in *C. elegans* is comparatively mature to that of more anatomically complex metazoans, the human genome encodes two gene products, SIDT1 and SIDT2, that have been purported to maintain a conserved role in the transport of extracellular dsRNA (23, 24, 25). Significantly, human SIDT2 was recently shown to be elevated in PD patient brains in conjunction with increased alpha-synuclein levels, and also colocalized to Lewy bodies (26). Thus, the functional duality represented by TNK2/SID-3 axis in the coordination of both dopamine and dsRNA import into the dopaminergic neurons of *C. elegans* warrants further mechanistic investigation. We previously established *C. elegans* as a genetic animal model with which to investigate and reveal numerous conserved genes and cellular pathways underlying progressive dopaminergic neurodegeneration, often predating their association to PD in humans (9, 27-37). A distinct advantage of this model is the capacity to rapidly quantify dopaminergic neurodegeneration with precision, at the single-neuron level, and with rigor across large isogenic populations of animals per experiment (38, 39). As with mammalian PD toxin models, the uptake of dopamine through DAT-1, the *C. elegans* homolog of human DAT, can be measured indirectly through exposure to the selective neurotoxin, 6-hydroxydopamine (6-OHDA) (40). Similarly, the transport of dsRNAs by *C. elegans* SID-1, which is also localized on the plasma membrane (18), can be functionally quantified by fluorescence in response to RNAi knockdown of GFP. Through the application of these discrete bioassays, we describe a previously unreported convergence of cellular mechanisms that impact dopaminergic neuron survival. Furthermore, we demonstrate that specific PD patient-associated SNPs in *TNK2* exert an analogous impact on dopaminergic neurodegeneration when introduced into the nematode counterpart, *sid-3*, by CRISPR gene editing. Taken together, the findings presented here illustrate a functional nexus whereby genetic mutations that influence dopamine levels in neurons intersect with the regulation of dsRNAs, key organismal modulators of epigenetic response, to modulate resilience or susceptibility to neurodegeneration.

## Results

### SID-3 modulates 6-hydroxydopamine neurotoxicity in *C. elegans*

Mammalian TNK2 has a regulatory function that serves to control dopamine synaptic and pre-synaptic dopamine levels by actively preventing clathrin-dependent endocytosis of the dopamine transporter, DAT (4). To determine if the *C. elegans* TNK2 ortholog, SID-3, similarly prohibits the endocytosis of the conserved worm dopamine transporter homolog, DAT-1, we treated *sid-3(ok973)* loss-of-function mutants (a 1330bp deletion allele) with 6-OHDA. Prior to 6-OHDA treatment, *sid-3(ok973)* worms were crossed into the wildtype (N2) background containing a chromosomally integrated transgene that enables GFP expression exclusively in dopamine neurons via the *dat-1* promoter [BY250; P_*dat-1*_::GFP (*vtIs7)*] for visualization of neurodegeneration. If SID-3 and TNK2 share conserved roles in regulation of dopamine uptake through DAT/DAT-1, there should be a reduction in the uptake of 6-OHDA in *sid-3(ok973)* mutants, resulting in decreased neurodegeneration. Indeed, when compared to GFP-only control animals, the *sid-3(ok973)* mutants displayed robust protection from 6-OHDA, even at the highest concentration (50mM), indicating an attenuation of dopamine uptake in the *sid-3(ok973)* mutant background, and evidence of functional homology between the human and worm proteins in the modulation of DAT-1 by endocytosis (Figure 1A,1C-F).

**Figure 1.**
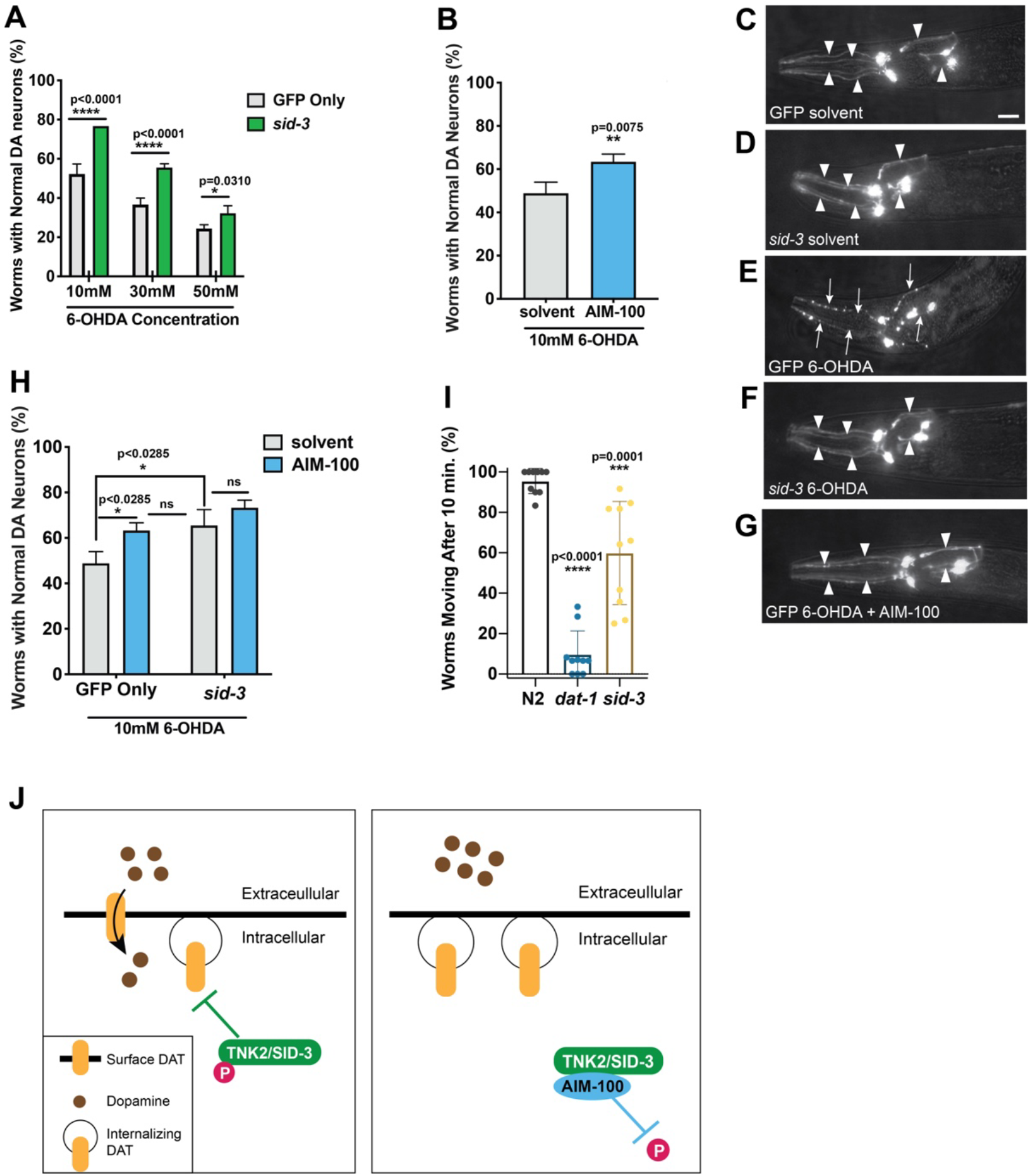
Inhibition of SID-3 protects *C. elegans* dopaminergic neurons from 6-OHDA toxicity. (A) Both wildtype (strain N2) *C. elegans* (grey bars) and *sid-3(ok973)* loss-of-function mutants with dopaminergic GFP expression (green bars) were exposed to 6-OHDA. Bars represent mean values of N=3. Error bars indicate SD; n= 30 worms in triplicate; two-way ANOVA with Sidak’s multiple comparisons. (B) Wildtype control (N2) animals with only GFP expressed in dopaminergic neurons were treated with either 1% ethanol solvent (grey bar) or 100µM AIM-100 (blue bar) and then subsequently assayed for neurodegeneration following treatment with 10mM 6-OHDA. Bars represent mean values of N=3. Error bars indicate SD with n=30 worms in triplicate; Student’s one-tailed *t*-test. (C-G) Neuron images labeled to reflect specified conditions; arrowheads and arrows indicate intact or degenerated dopaminergic neurons, respectively. Scale bar, 20 μm. (H) GFP-only control and *sid-3(ok973)* mutant animals treated with either 1% ethanol solvent (grey bars) or 100µM AIM-100 (blue bars) and then assayed for neurodegeneration following exposure to 10mM 6-OHDA. Bars represent mean values of N=3. Error bars indicate SD with n=30 worms in triplicate; two-way ANOVA with Tukey’s post hoc multiple comparisons test. (I) Wildtype (N2) worms, and *dat-1(ok157)* or *sid-3(ok973)* deletion mutants were assayed for dopamine-specific behavioral change by swimming-induced paralysis (SWIP). Bars represent mean values of N=10. Error bars indicate SD, with 10-15 animals scored per replicate; one-way ANOVA with Dunnett’s post hoc multiple comparisons test. (J) Schematic model of TNK2/SID-3 regulation of DAT endocytosis. Left panel, baseline/wildtype conditions with activated TNK2/SID-3 blocking transporter endocytosis, thereby enabling 6-OHDA uptake and neurodegeneration; Right panel, AIM-100 inhibition of TNK2/SID-3 phosphorylation results in diminished endocytic activity and protection from 6-OHDA neurotoxicity. TNK2/SID-3 (green); phosphate (pink); DAT (yellow); dopamine (brown); AIM-100 (blue).

To support our findings implicating SID-3 in the regulation of DAT-1 endocytosis, we treated wildtype (GFP-only) animals with an established and selective TNK2 inhibitor, AIM-100 (41), that has been shown to inhibit the Tyr284-phosphorylation of TNK2 (42) and prevent the block it maintains on the endocytosis of mammalian DAT (4). Animals were treated daily with AIM-100 for one-hour, while control groups received only treatments with the solvent (1% ethanol). Like the *sid-3(ok973)* mutants, GFP-only animals treated with AIM-100 display significant protection from the neurotoxicity of 6-OHDA when compared to solvent controls (Figure 1B,1G). Additionally, we treated *sid-3(ok973)* deletion mutants with AIM-100 to determine if the increase in neuroprotection observed in wildtype animals would be indicative of the drug possibly working through a target distinct from SID-3. AIM-100 did not enhance the neuroprotection of *sid-3(ok973)* mutants when compared to the solvent-only treated *sid-3(ok973)* mutants (Figure 1H). Moreover, solvent-only treated *sid-3(ok973)* mutants were not significantly different from AIM-100-treated (GFP-only) animals (Figure 1H). Considering the equivalence in neuroprotection observed with AIM-100-treated wildtype animals and untreated *sid-3(ok973)* mutants, and that this protection is abolished in a *sid-3 (ok973)* mutant background, it is reasonable to conclude that SID-3 is the target of AIM-100 in *C. elegans*, and that this drug provides neuroprotection by attenuating 6-OHDA uptake into the dopaminergic neurons of *C. elegans*.

To further demonstrate the impact of SID-3 on dopamine uptake in *C. elegans*, we conducted a behavioral analysis where we monitored *sid-3(ok973)* mutants in swimming-induced paralysis (SWIP) assays (43). SWIP assays are used as an indirect measure of proper synaptic dopamine clearance. Animals that display significantly aberrant dopamine clearance undergo paralysis after 10-minutes of thrashing in water, while animals with intact dopamine clearance can thrash continuously (43). Our results, as depicted in Figure 1I, demonstrate that *sid-3(ok973)* animals exhibited significantly more paralysis than did wildtype (N2) controls after 10-minutes of thrashing. However, this was not as significant as *dat-1 (ok157)* loss-of-function mutants (Figure 1I). It is not surprising that *sid-3(ok973)* mutants do not have as pronounced of an effect as the *dat-1* deletion mutants since a limited pool of DAT would remain at the presynaptic plasma membrane, albeit undergoing an increase in basal internalization (44).

Collectively, these results, using both genetic and pharmacological modifiers, suggest that SID-3 retains a conserved function of mammalian TNK2 in preventing the endocytic internalization of DAT (Figure 1J). This model predicts that either deletion [*sid-3(ok973)*] or inhibition [by AIM-100] of SID-3/TNK2 activity would be neuroprotective, with a net increase of synaptic dopamine resulting in enhanced dopaminergic neurotransmission. Importantly, since this effect would presumably be lost when *TNK2* is mutant, these results therefore suggested to us that the SNPs in *TNK2*, inherited recessively by PD patients, were potentially representative of *increased* TNK2 stability and/or activity.

### NAB2-induced neuroprotection requires the *C. elegans* NEDD4 homolog, WWP-1

Since a constitutive blockade of DAT endocytosis by TNK2 would rapidly deplete the levels of synaptic dopamine, it is critical that the dopaminergic system be both dynamic and responsive to a variety of conditions. NEDD4 is a E3 ubiquitin ligase known to ubiquitinate TNK2 for degradation (6, 45). To determine if increasing the activity of the NEDD4 homolog in *C. elegans*, WWP-1, would lead to neuroprotection from 6-OHDA, we treated wildtype (GFP-only) animals with a previously defined and selective NEDD4 activator, NAB2 (9). If NEDD4 has a regulatory effect on DAT internalization by targeting TNK2/SID-3 for degradation, then an increase in NEDD4/WWP-1 activity would result in a reduction of 6-OHDA import, and neuroprotection. Indeed, when compared to solvent-treated controls, NAB2-treated animals were protected from 6-OHDA-induced neurodegeneration (Figure 2A), in accordance with prior work where we demonstrated that alpha-synuclein-induced toxicity was attenuated by NAB2 treatment in transgenic *C. elegans*, complementing supporting results in both yeast and rat primary neurons (9). It should be noted that, because the *C. elegans* genome lacks an endogenous alpha-synuclein gene, our new findings acquired in the complete absence of alpha-synuclein are mechanistically significant, since alpha-synuclein was reported to be a direct target of NEDD4-dependent ubiquitination and degraded through endolysosomal clearance (46). Taken together, the regulatory node centered at NEDD4/TNK-2 functionality promotes neuroprotection through multiple means.

**Figure 2.**
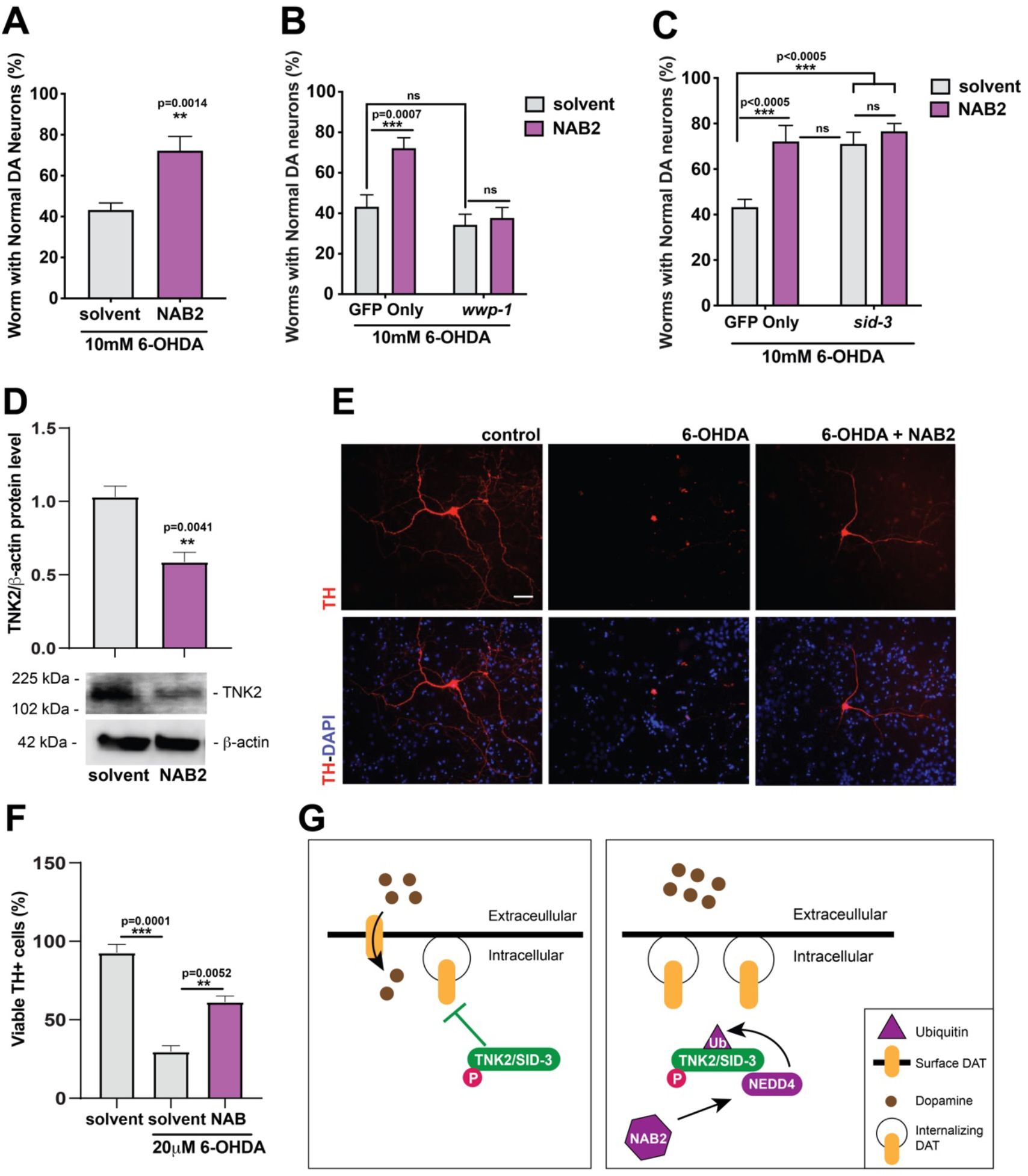
Rat primary neurons display decreased TNK2 levels and dopaminergic neuroprotection from 6-OHDA in response to NEDD4/WWP1 activation by NAB2 treatment. (A-C) Specified worm strains treated with 100µM NAB2 (purple bars) or 1% ethanol solvent (grey bars) received a 1-hour treatment each day before exposure to 10mM 6-OHDA. (A) Wildtype GFP-only control animals, where bars represent mean values of N=3. Error bars indicate SD with n= 30 worms in triplicate; Student’s one-tailed *t*-test. (B) GFP-only and *wwp-1(gk372)* loss-of-function animals, where bars represent mean values of N=3. Error bars indicate SD with n=30 worms in triplicate; two-way ANOVA with Tukey’s multiple comparisons. (C) GFP-only and *sid-3(ok973)* animals, where bars represent mean values of N=3. Error bars indicate SD with n=30 worms in triplicate; two-way ANOVA with Tukey’s multiple comparisons. (D) Rat cortical neurons were treated with 2µM NAB2 or DMSO solvent and western blotting was used to detect TNK2. The representative blot shown here, with the accompanying graph representing the mean of four experiments, indicate a significant decrease in TNK2 levels in response to NAB2. Error bars indicate SEM; Student’s two-tailed *t*-test. (E, F) Rat dopaminergic neurons were treated with NAB2 or the solvent for NAB2 (DMSO) and 20 mM 6-OHDA. Tyrosine hydroxylase positive (TH+) neurons were assessed by immunocytochemistry. NAB2 treatment prevents the dopamine neuron degeneration induced by 6-OHDA (E) Representative neurons; Scale bar 50µm. Red: TH+ cells; Blue: DAPI+ cells. (F) Quantitative analysis of viable TH+ dopaminergic cells where bars represent mean values of N=3, error bars indicate SEM with 20-30 micrographs counted per experiment: one-way ANOVA with Tukey’s multiple comparisons. (G) Schematic diagrams depicting regulation of DAT endocytosis and dopamine/6-OHDA uptake by TNK2/SID-3 (left panel); ubiquitination of TNK2/SID-3 in response to NAB2 activation of NEDD4/WWP-1 (right panel) reduces TNK2/SID-3 levels and enhances dopaminergic neuroprotection. TNK2/SID-3 (green); phosphate (pink); DAT (yellow); dopamine (brown); NEDD4 (purple); NAB2 (purple hexagon); Ubiquitin (purple triangle).

Although *C. elegans* WWP-1 shares the highest overall sequence homology and conserved domain structure with human NEDD4, and *wwp-1* is highly expressed in the nervous system, the worm genome contains multiple E3 ligases that share varying degrees of homology with NEDD4. Therefore, to confirm that the observed responses to NAB2 were a consequence of its action on WWP-1, we crossed animals with a WWP-1 loss-of-function mutant, *wwp-1(gk372*), into GFP-only animals and assayed these with 6-OHDA and NAB2 treatments. Animals in the *wwp-1(gk372)* mutant background showed a complete abolishment of NAB2-induced neuroprotection from 6-OHDA, while control animals retained NAB2-neuroprotection (Figure 2B), thereby demonstrating that NAB2 acts through the NEDD4 homolog, WWP-1, in *C. elegans*. We also note that the *wwp-1(gk372)* mutation did not enhance neurodegeneration independently (Figure 2B). We also wanted to examine if NAB2-neuroprotection from 6-OHDA operates through the targeting of SID-3 by WWP-1 for degradation, consequentially enabling DAT-1 internalization. As depicted in Figure 2C, treatment of *sid-3(ok973)* mutants either with NAB2 or its solvent control did not affect the dopaminergic neuroprotection that was displayed in the absence of SID-3 function (Figure 2C). Furthermore, NAB2 treatment of wildtype (GFP-only) worms offered a similar level of protectidon from 6-OHDA neurotoxicity as was observed with *sid-3(ok973)* mutant controls (Figure 2C). Thus, in terms of neuroprotection, the activation of NEDD4/WWP-1 by NAB2 phenocopies the complete depletion of TNK2/SID-3 in *C. elegans*.

### Activation of NEDD4 in rat primary neurons leads to a reduction of endogenous TNK2 and neuroprotection from 6-OHDA

To determine if the proposed mechanism of dopaminergic regulation we have elucidated in *C. elegans* translates to mammals, we examined rat cortical primary neurons for evidence of a change in TNK2 levels following activation of NEDD4 by NAB2. We previously reported that rat primary neurons exhibited neuroprotection in response to NAB2 treatment (9). Indeed, this effect further translated to human neurons derived from iPSCs of PD patients (8). It remains to be discerned if the neuroprotective effects observed are a potential consequence of a decrease in TNK2 levels. We demonstrated that enhancement of NEDD4 activity by NAB2 treatment lowers endogenous neuronal TNK2 protein levels (Figure 2D). Similar to the worm experiments described in 2A, we wanted to determine if increasing the activity of NEDD4 in rat primary dopaminergic neurons would lead to neuroprotection from 6-OHDA. Cultured neurons were treated to NAB2, with or without 6-OHDA for 24 hours (Figure 2E, F). The control group was treated with the solvent only. Compared to solvent-treated controls, there were more TH+ viable neurons in the NAB2-treated group (Figures 2E,F), consistent with the neuroprotection observed in *C. elegans* (Figure 2A). The combined outcomes of these studies can be summarized by a model (Figure 2G) that portrays both the targeting of TNK2 for degradation via the ubiquitin-proteasome system and altered endocytic regulation as potential cellular mechanisms amenable to fine-tuning of dopamine levels to sustain optimal dopaminergic neurotransmission and health.

### SID-3 facilitates dsRNA-mediated gene silencing in *C. elegans* dopaminergic neurons

SID-1-dependent transport of dsRNA in *C. elegans* is modulated through the internalization of this plasma membrane protein by clathrin-dependent endocytosis (22). Previous studies reported that SID-3 plays a functional role in support of the systemic spread of dsRNA between the somatic cells of *C. elegans*, with a concomitant impact on the efficacy of RNAi (18). Just as TNK2 regulates the clathrin-dependent endocytosis of DAT to modulate dopamine reuptake, it has been hypothesized that SID-3 functions as an endocytic modulator of dsRNA uptake by preventing the clathrin-dependent endocytosis of SID-1 (18). To determine if SID-3 regulates dopaminergic RNAi, we fed animals RNAi-inducing bacteria engineered to produce exogenous dsRNA targeting GFP that was expressed exclusively in the dopaminergic neurons. Due to an inherently RNAi resistant nature of neurons in *C. elegans* (47), GFP-only and *sid-3(ok973)* animals were crossed into an RNAi-hypersensitive background *eri-1(mg366)*, which does not impact the transport of dsRNAs but rather has changes in exonuclease and nucleic-acid-binding activity that facilitate RNAi efficacy (48). Subsequent RNAi knockdown in wildtype animals led to silencing of GFP fluorescence, when compared to empty vector (EV) RNAi control (Figures 3A, C, D). In contrast, *sid-3(ok973)* mutants displayed significantly less dopaminergic neuron silencing compared to wildtype animals (Figure 3A, E). As has been previously reported for *sid-3* mutants in other cell types (18), a loss of SID-3 activity diminished, but did not completely abolish, RNAi silencing; an expected result indicative of increased endocytosis of SID-1 at the plasma membrane, where a modicum of this dsRNA transporter likely remains present (Figure 3G). We further demonstrated the necessity of active SID-3 for dsRNA-induced silencing in the dopaminergic neurons by treating animals with AIM-100. Wildtype worms treated with AIM-100 exhibited a decrease in dopaminergic neuron fluorescence in response to RNAi knockdown of GFP when compared to solvent-only controls (Figure 3B, F). This result was consistent with our prior data demonstrating that both AIM-100 treatment of wildtype animals and *sid-3(ok973)* mutants similarly displayed neuroprotection from 6-OHDA (Figure 1A, B). In this case, however, lifting the block on dsRNA uptake as a response to AIM-100 inhibition of SID-3 results in an increased capacity to respond to dsRNA that is readily observed through the efficacy of gene silencing (Figure 2G).

**Figure 3.**
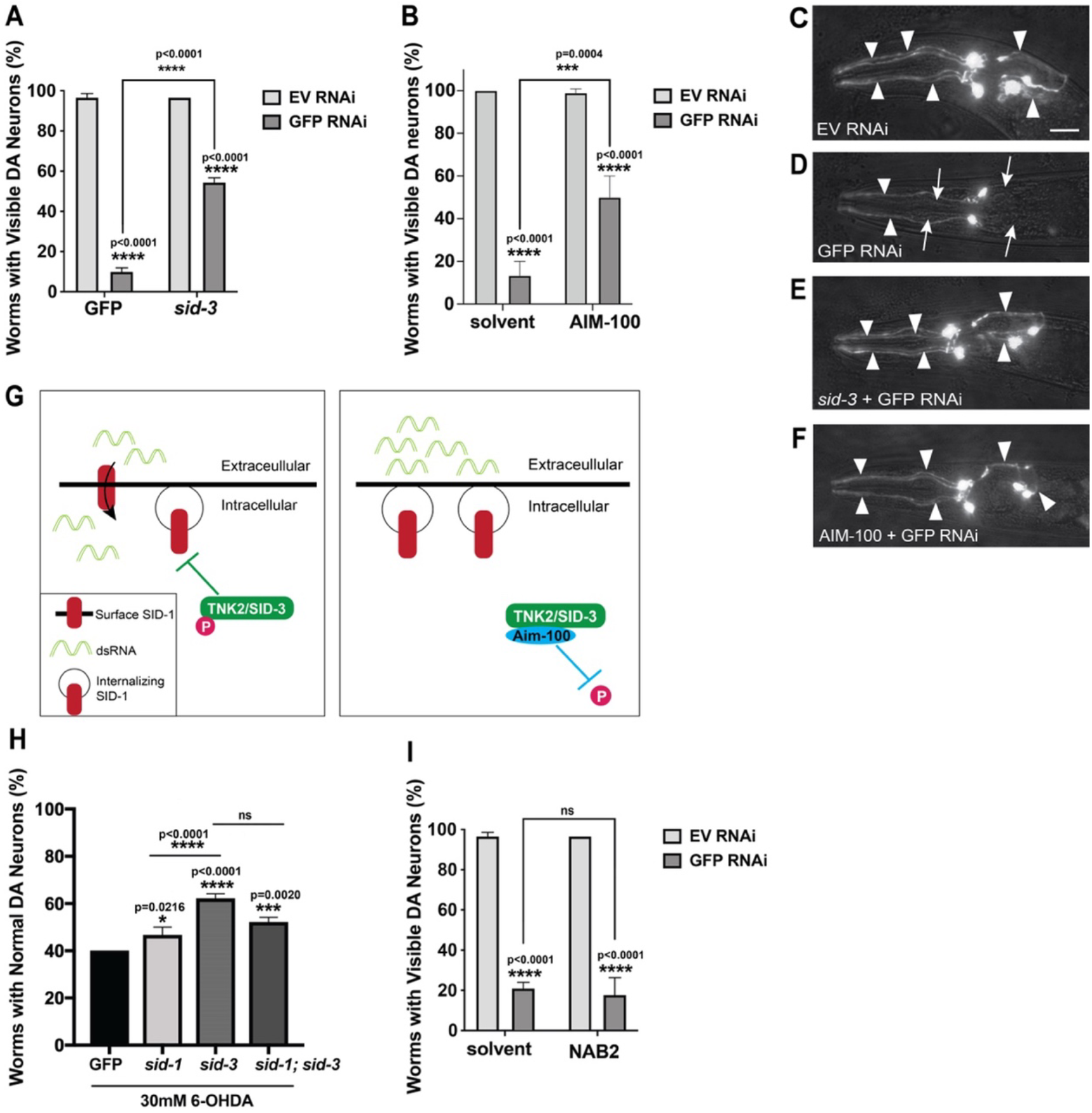
Inhibition of SID-3 decreases dopaminergic RNAi efficacy. (A) Animals expressing GFP in the dopaminergic neurons of both control (wildtype; GFP-only) and *sid-3(ok973)* mutant animals in an RNAi-hypersensitive background [*eri-1(mg366)*] fed bacteria harboring dsRNA targeting GFP (dark grey bars) or empty vector (EV; light grey bars). Bars represent mean values of N=3. Error bars indicate SD with n=30 worms in triplicate; two-way ANOVA with Tukey’s post hoc multiple comparisons test. (B) GFP-only animals in a RNAi-hypersensitive background raised on RNAi plates with or without 100µM AIM-100 added to the agar. Bars represent mean values of N=3. Error bars indicate SD with n=30 worms in triplicate; two-way ANOVA with Tukey’s post hoc multiple comparisons test. (C-F) Representative dopaminergic neuron images of worms from experiments in panels A-B where neurons are either still visible (arrowheads) or GFP is silenced (arrows). Scale bar, 20 μm. (G) Schematic diagram of TNK2/SID-3 regulation of dsRNA import. Left panel depicts activated TNK2/SID-3 preventing the endocytosis of SID-1, thereby facilitating dsRNA transport; right panel, inhibition of TNK2/SID-3 activity by AIM-100 releases the block on endocytosis, causing resistance to dsRNA transport. TNK2/SID-3 (green); phosphate (pink); SID-1 (red); dsRNA (light green); AIM-100 (blue). (H) Direct comparison of wildtype control (GFP-only), *sid-1(pk3321), sid-3(ok973)*, or *sid-1(pk3321);sid-3(ok973)* double mutants when exposed to 30mM 6-OHDA. Distinctions potentially reflect the relative contributions to neuroprotection from dsRNA uptake and/or dopamine (6-OHDA) transport. Bars represent mean values of N=3. Error bars indicate SD with n= 30 worms in triplicate; one-way ANOVA with Tukey’s multiple comparisons. (I) GFP-only animals in a RNAi-hypersensitive background raised on RNAi plates with or without NAB2 (100µM) added to the agar do not exhibit a significant change in RNAi efficacy. Bars represent mean values of N=3. Error bars indicate SD with n=30 worms in triplicate; two-way ANOVA with Tukey’s post hoc multiple comparisons test.

In considering the robust neuroprotection observed with *sid-3(ok973)* mutants, we reasoned that the effect could be partially attributed to the diminished import of silencing dsRNAs. To test this hypothesis, we assayed *sid-1(pk3321)* loss-of-function mutants with 6-OHDA and compared the results to *sid-3(ok973)* or *sid-1(pk3321)*; *sid-3(ok973)* double mutants, as well as wildtype (GFP-only) control animals (Figure 3H). Although depletion of the SID-1 dsRNA transporter in *sid-1(pk3321)* mutants was neuroprotective against 6-OHDA, compared to GFP only control animals, it was not as nearly protective as observed in the *sid-3(ok973)* deletion mutants. Likewise, *sid-1(pk3321)*; *sid-3(ok973)* double mutants also displayed significantly increased levels of neuroprotection, similar to *sid-3 (ok973)* alone (Figure 3H). Superficially, these data could be suggestive of an epistatic relationship between the *sid-1* and *sid-3* alleles. Yet, in the specific case of the dopaminergic system, the enhanced neuroprotection exhibited in *sid-3(ok973)*, along with evidence of intersecting regulation of dopamine and dsRNA uptake by SID-3, suggest that a more complex regulatory dynamic is at work in response to the acute neurodegenerative challenge presented by 6-OHDA treatment.

In contrast to what was observed using AIM-100 (Figure 3B,F), a distinction with NAB2 treatment was observed, as this NEDD4/WWP-1 activator did not alter the extent of dopaminergic GFP RNAi silencing (Figure 3I). We speculate that SID-3 may bind to DAT and SID-1 through alternative motifs and, when it is bound to SID-1, the activated NEDD4/WWP-1 E3 ligase is unable to interact with TNK2/SID-3 to target it for degradation. This is supported by evidence that the NCK-1/SID-4 protein, a SH2/SH3 domain-containing adaptor protein, is also required for SID-3-dependent RNAi silencing (49). A putative complex between SID-3 and SID-4 may represent a steric hinderance to NEDD4/WWP-1 interaction. The observed specificity of NAB2 as an effector of only the dopaminergic regulatory contribution to the neuroprotection observed illustrates a subtle but significant difference in the endocytic regulation of dopamine compared to dsRNA transport. The chemical-genetic approach employed here highlights the utility of *C. elegans* models in parsing out the intricacies of cellular mechanisms (50). This strategy represents an effective means for more rapid, cost-effective, and functionally justified prioritization of therapeutic targets for preclinical research in neurodegenerative diseases (51).

### Consequences of PD-patient associated SNPs on TNK2/SID-3 function *in vivo*

*TNK2* encodes a non-receptor tyrosine kinase with a unique domain structure that is conserved in *C. elegans* SID-3 (Figure 4A). Mutations in *TNK2* have been identified in several types of cancers (52). Among protein regions that are mutated are the protein kinase domain (PKD), a proline rich NEDD4 interacting region (P), and ubiquitin associating domain (UBA). Previous research focused on infantile-onset epilepsy showed that a similar mutation of *TNK2* in the proline-rich region blocked its interaction with NEDD4, eventually leading to infant mortality (53). Of particular interest to our study, SNPs within *TNK2* discovered through whole exome sequencing of DNA from patients with familial PD include variants in some of these predicted functional domains (1) (Figure 4A).

**Figure 4.**
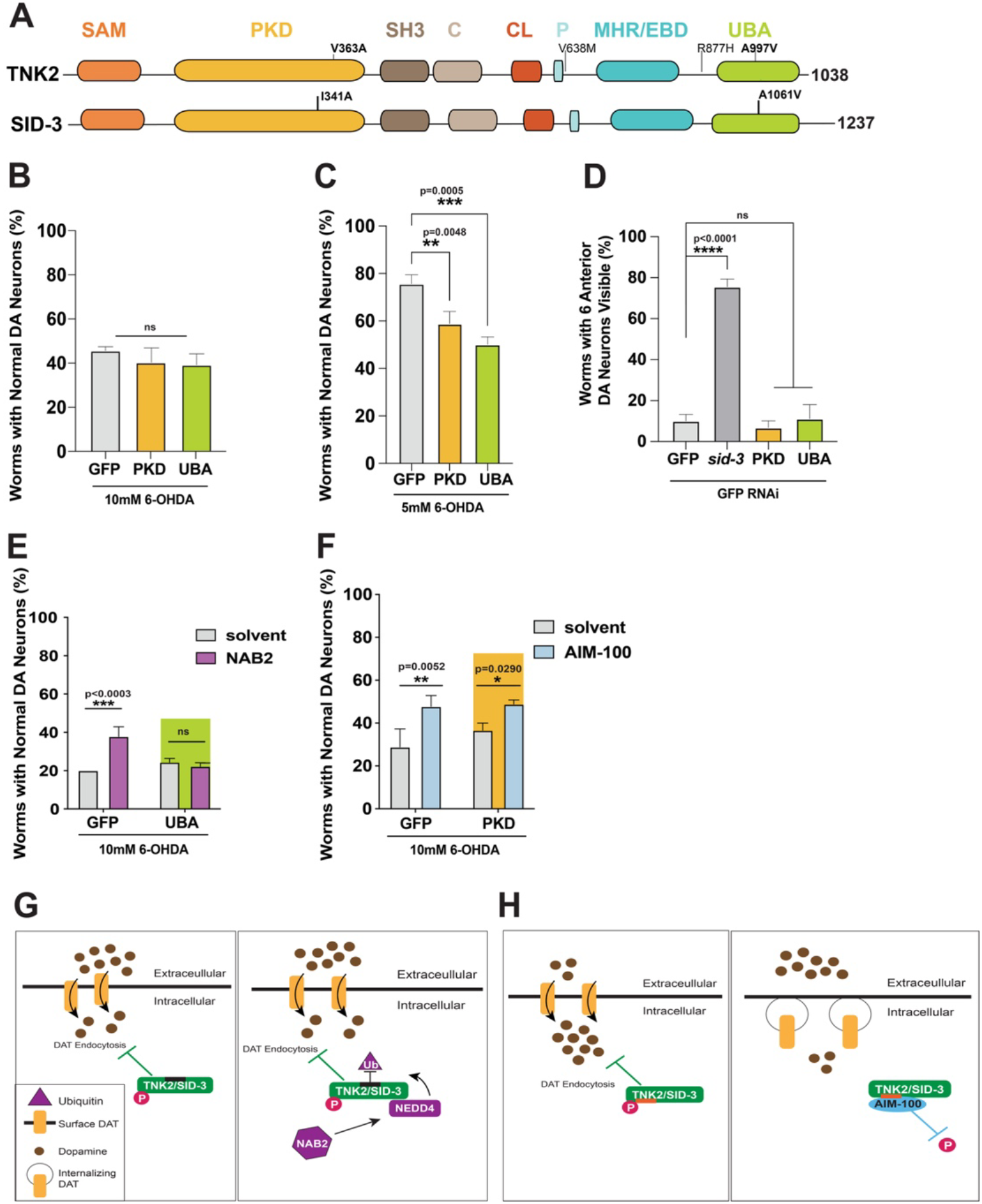
Functional effects of PD-related SNPs in *TNK2* at analogous positions in *sid-3* of CRISPR-edited *C. elegans*. (A) Protein domains of human TNK2 share homology with SID-3. SAM = sterile alpha motif; PKD = tyrosine protein kinase domain; SH3 = Src homology domain; C = CDC42-binding domain; CL = clathrin-interaction domain; P = proline rich motif containing the NEDD4 binding site; MHR/EBD = MIG6 homology region (also known as an Epidermal growth factor receptor Binding Domain or EBD); UBA = ubiquitin association domain. Amino acid changes resulting from the PD-related SNPs in human *TNK2* are indicated in black; the CRISPR-edited sites in the corresponding SID-3 protein of *C. elegans* are indicated, with the human SNPs bolded. (B) Wildtype control (GFP-only) or protein kinase domain (PKD) mutant SID-3(I341A) or ubiquitin association-domain (UBA-domain) mutant SID-3(A1016V) CRISPR-edited animals exposed to either 10mM 6-OHDA or (C) 5mM 6-OHDA. (D) RNAi-hypersensitive worms in either the GFP-only (WT SID-3), *sid-3(ok973)* deletion mutant, or CRISPR-edited backgrounds (with PKD or UBA SNPs) following RNAi feeding with bacteria targeting GFP for two generations. (B-D) Bars represent mean values of N=3. Error bars indicate SD with n=30 worms in triplicate; one-way ANOVA with Dunnett’s post hoc multiple comparisons test. (E) Animals with the UBA-domain SNP treated with 100µM NAB2 are not protected from 6-OHDA-induced neurodegeneration. Bars represent mean values of N=3. Error bars indicate SD with n=30 worms in triplicate; two-way ANOVA with Sidak’s post hoc multiple comparisons test. (F) Animals with the PKD SNP treated with 100µM AIM-100 are still protected from 6-OHDA. Bars represent mean values of N=3. Error bars indicate SD with n=30 worms in triplicate; two-way ANOVA with Sidak’s post hoc multiple comparisons test. (G) SID-3 containing the PD-patient derived SNP in the UBA domain at an analogous position that found in TNK2 is depicted as being targeted for ubiquitin-mediated degradation without (left panel) and with (right panel) NAB2-activated NEDD4 E3 ligase activity, which is blocked by the UBA-domain SNP (H) SID-3 engineered to contain the PD-patient derived SNP found in the PKD of TNK2 is depicted as hyperactive in preventing DAT endocytosis, thereby resulting in depletion of synaptic dopamine (left panel); this SNP does not impact the ability of SID-3 to be inactivated by AIM-100, and therefore retains the capacity for DAT internalization and neuroprotection (right panel). Mutated TNK2/SID-3 (green with dark purple bar representing UBA-SNP); or (green with orange bar representing PKD-SNP); phosphate (pink); DAT (yellow); dopamine (brown); NEDD4 (purple); NAB2 (purple hexagon); Ubiquitin (purple triangle).

We sought to determine if two of the PD patient-associated SNPs located in conserved sites of post-translational regulation of TNK2/SID-3 activity (PKD) or stability (UBA domain) confer an increased susceptibility to neuronal dysfunction and/or survival. To examine these SNPs *in vivo*, we used CRISPR technology to edit the chromosomal *C. elegans sid-3* gene and thereby generate two separate new strains encoding SNPs in either the conserved PKD or UBA domains of the endogenous gene product that reflected the mutations found in PD patients (1) (Figure 4A). We first tested if the modified nematodes containing these specific SNPs exhibited an enhanced susceptibility to 6-OHDA-induced neurodegeneration. However, no increase in dopaminergic neurodegeneration was observed in animals using the standard assay conditions for exposure, 10mM 6-OHDA (Figure 4B). We reasoned that the acute neurotoxicity of 10mM 6-OHDA could mask the impact of either genetic background. Therefore, we decided to half the dosage to provide for a more sensitive test. Exposure to 5mM 6-OHDA revealed that the SNP-containing worms showed significantly more neurodegeneration and that these animals were more sensitive to the lower dosage of neurotoxin than their WT counterparts (Figure 4C). These results provide mechanistic evidence for these specific PD-patient SNPs in TNK2/SID-3 functionally altering dopamine uptake, via its neurotoxic proxy molecule, 6-OHDA.

We next wanted to evaluate these same PD-patient SNPs for their impact on dsRNA import to induce RNAi silencing in *C. elegans*. Therefore, both of the CRISPR-edited strains were crossed into the hypersensitive *eri-1(mg366)* background and fed RNAi bacteria selectively targeting GFP. Both the PKD and UBA domain-associated SNPs led to very pronounced GFP silencing in the dopaminergic neurons (Figure 4D). This was striking when compared to the resistance to RNAi observed in *sid-3(ok973)* loss-of-function mutant animals within the same the hypersensitive background. In fact, the level of RNAi knockdown observed in either of the CRISPR-edited strains was largely equivalent to the activity observed in non-mutant controls. Whereas an impact on dopaminergic neurodegeneration by SNPs associated with PD patients could have been expected, the added observation that introduction of these same polymorphisms into SID-3 also modulates the epigenetic machinery of RNAi silencing in the same neurons provides a new prospective through which to consider gene-by-environment interactions in PD.

To further investigate the functional consequences of these polymorphisms, we examined whether the NAB2-induced neuroprotection previously observed remained intact in animals encoding SID-3 with the UBA domain-associated SNP. Interestingly, NAB2 was no longer neuroprotective against 6-OHDA in these SNP-containing animals (Figure 4E). This indicates that the interaction between SID-3 and WWP-1 was abolished and is necessary for proper DAT-1 regulation (Figure 4G). These data are consistent with studies done in mammalian cell cultures on similar, but not PD-associated, *TNK2* variants harboring SNPs shown to disrupt NEDD4-TNK2 interactions (53). Therefore, failure of NEDD4-dependent degradation of TNK2 provides a mechanism to explain how patients with a recessive mutation in the *TNK2* UBA domain manifest with the symptoms of PD as a consequence of increased DAT-mediated reuptake of dopamine and a reduction in synaptic dopamine.

Treatment of *C. elegans* with the TNK2 inhibitor, AIM-100, leads to neuroprotection in a manner that phenocopies SID-3 loss-of-function [(*sid-3 (ok973)*] (Figure 1A, B). In contrast, when CRISPR-edited worms containing a SNP in the PKD of SID-3 were treated with AIM-100, neuroprotection persisted (Figure 4F). Notably, a similar PKD mutation in human TNK2 was found in prostate cancer that actually increased TNK2 activity (42). In the dopaminergic system, where we have demonstrated that depletion of this gene product is neuroprotective, a hyperactive TNK2/SID-3 would be predicted to exert detrimental consequences that could contribute to PD. Importantly, it was determined that AIM-100 still functioned to prevent the phosphorylation of TNK2 in the hyperactive variant (42), thereby correlating with our observation that AIM-100 retains its inhibitory/protective activity even when SID-3 contains an analogous SNP. Thus, in the absence of AIM-100, these same CRISPR-modified animals are no longer protected from neurodegeneration (Figure 4F,4H). These results define the clinical impact of polymorphisms in *TNK2*, identified by WES of PD patients, as an outcome of a lost cellular capacity to either degrade or inactivate this pivotal enzyme. These results exemplify an experimental strategy whereby the directed application of invertebrate model systems serves as a means to advance functional annotation of human genomic variation, inform therapeutic discovery, and accelerate clinical understanding (51, 54).

As depicted in Figure 5, these data collectively support a molecular model for dopaminergic neuroprotection that reflects how the neuromodulation of synaptic and intracellular dopamine levels directly converges with the organismal machinery required to transmit exogenous influences on gene expression conferred through non-coding RNAs. A prerequisite to an eventual understanding of gene x environmental contributions in discerning disease risk is a mechanistic means through which their combined effects are manifested. A failure of such integrated processes may underlie differential susceptibility to disease, and as demonstrated here, the neurodegeneration observed in individuals with PD.

**Figure 5.**
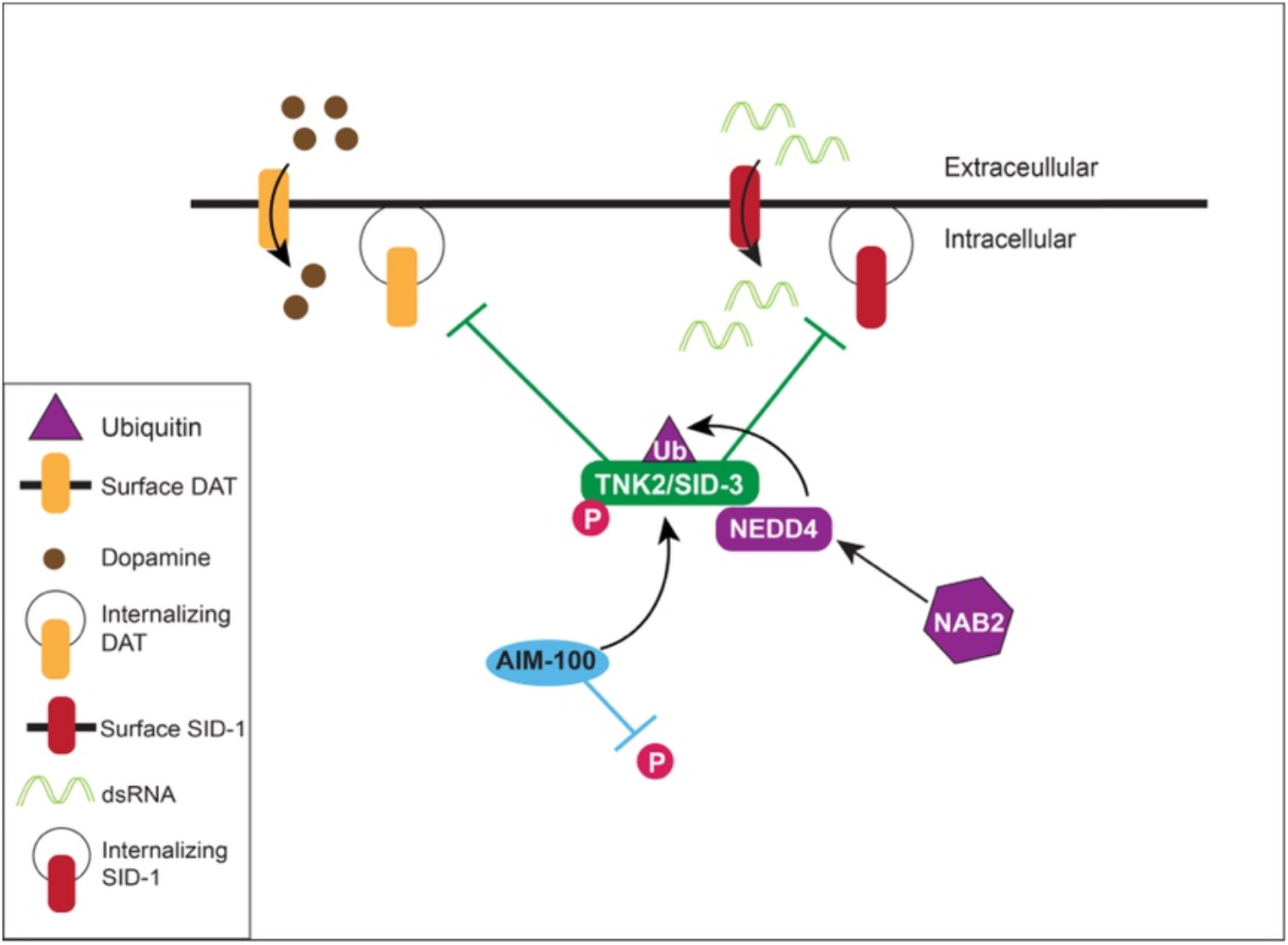
Model depicting the integrated dopaminergic and epigenetic modulation of neuroprotection through TNK2/SID-3. The endocytosis of the dopamine transporter, DAT, is necessary for maintaining levels of synaptic and cytoplasmic dopamine in a healthy balance that ensures proper neurotransmission. The capacity of the dopaminergic system to adapt to an imbalance in this system imposed by a variety of stressors. As shown above, the non-receptor tyrosine-kinase, TNK2, maintains a functional hold on the endocytosis of the dopamine transporter, DAT, in the regulation of dopamine levels. NEDD4, an evolutionarily conserved E3 ubiquitin ligase, interacts with TNK2, and targets it for degradation by the ubiquitin-proteasome system (UPS). We demonstrated that NEDD4 activation, which is induced by treatment with the small molecule, NAB2, attenuates dopaminergic neurodegeneration induced by the neurotoxic dopamine analog, 6-OHDA, in both in *C. elegans* and rat primary neuron cultures. In an analogous manner, the *C. elegans* TNK2 ortholog, SID-3, is known to be required for epigenetic gene silencing by RNAi, as a result of its function in systemic RNAi through the regulation of the SID-1 dsRNA transporter protein. Our results show that either genomic deletion or chemical (AIM-100) inhibition of SID-3 abolished dsRNA-targeted silencing of gene expression in worm dopamine neurons – in addition to protecting both *C. elegans* and rat primary dopamine neurons from 6-OHDA neurotoxicity. We used *C. elegans* to functionally examine why this protective capacity is lost in PD patients that contain SNPs in *TNK2*. By engineering SID-3 in worms to mimic specific PD-patient SNPs found in*TNK2*, we have revealed that the mechanism of neurodegeneration in these patients is likely not due to a loss-of-function but is instead a consequence of sustained TNK2/SID-3 activity that affects both dopamine and dsRNA-mediated gene silencing in *C. elegans*, and possibly humans. This model, and the collective outcomes of our results, represent a functional convergence and integrated control of dopaminergic and epigenetic mechanisms that have direct implications for modulating neurodegeneration in PD, and possibly other dopamine-related disorders. TNK2/SID-3 (green); phosphate (pink); DAT (orange); dopamine (brown); AIM-100 (blue); SID-1 (red); dsRNA (light green); NEDD4 (purple); NAB2 (purple hexagon); Ubiquitin (purple triangle).

## Discussion

The symptomatic phenotypes of PD are largely a result of the death of dopaminergic neurons located in the *substantia nigra*. The clinical emergence of symptoms, such as resting tremor and rigidity, arise only once 50-80% of the dopaminergic neurons are lost yet the age of onset can vary dramatically among individuals (55). The search to uncover new genetic associations to PD is ongoing and the significance of distinct cellular pathways that impact the dopaminergic system, and the intersection of them, will be critical to defining prognosis and closing the gap between diagnosis and treatment (56). As human genomic data flood the present research landscape with tantalizing associations and clues to disease mechanisms, the challenging task to functionally annotate these genomic variants has become increasingly significant. Research consortia have arisen to tackle the challenges of rare and undiagnosed diseases (57, 58, 59), joining the efforts of individual labs in parsing the data deluge with experimentation to accelerate therapeutic target and drug discovery for more common diseases as well (37, 60-63). These collective efforts have begun to bear fruit, and they are not limited to the direct relationship between genotype and phenotype alone. As the present study strives to impart, experimental strategies inclusive of the complexities introduced by epigenetic regulation, layered on a background of disease-modifying factors, represent a next level of functional genomic analysis (64, 65).

Fundamental organismal strategies as drivers of evolutionary change typically manifest as an increased capacity to adapt to fluctuating environments. This is epitomized by the epigenetic control of gene expression (66). Medical research has merely scratched the surface of what must urgently be understood to attain a clearer picture of how external (environmental) epigenetic influences on organismal health intersect with heritable or *de novo* genetic susceptibilities to manifest in disease states. A quantitative measure, termed stochastic epigenetic mutation (SEM), was devised to represent dysregulation at sites of genomic DNA methylation, an epigenetic hallmark that increases with chronological age (67). More recently, another metric, epigenetic mutation load (EML), has been applied by clinical researchers to the analysis of PD risk (15). EML represents the aggregate value of SEMs within the genome of an individual and is intended to reflect overall change or failure of epigenetic maintenance systems. Application of EML analysis to patient data on PD-associated loci revealed genes at greater risk for being affected by epigenetic modulation. Among a short list of only five such genes identified as having the strongest correlation to epigenetic modulation in PD patients was *NEDD4* (15).

Whereas TNK2 has been heavily studied in the field of cancer (52), it was only more recently identified as mutated in patients with PD (1). We therefore attempted to discern the mechanistic significance of TNK2/SID-3 activity in PD by addressing three key points: 1) we determined that SID-3 retains the conserved role of TNK2 in modulating reuptake of dopamine in *C. elegans*; 2) we found that SID-3 also regulates the capacity for dsRNA-induced gene silencing in the dopaminergic neurons and that this modulates neurodegeneration; and 3) we evaluated the functional consequences of PD patient-associated SNPs in TNK2 using SID-3 for their respective impact on neurodegeneration and dsRNA-mediated gene silencing *in vivo*. Our findings indicate that the mutant variants of *TNK2* found in patients define a molecular etiology for PD in which neurodegeneration is a consequence of dysfunctional cellular dynamics at an intersection of dopaminergic and epigenetic signaling.

The core cellular machinery for dopamine biosynthesis, vesicular packaging, transport, and neurotransmission is highly conserved among metazoans, as is the tight regulation of its component proteins. Invertebrate species, most prominently *Drosophila melanogaster* and *C. elegans*, have provided numerous insights into mechanistic features of dopaminergic neurotransmission and neurodegeneration (68-70). Although the genes for key enzymes and proteins (i.e., *DICER1, DROSHA1*, Argonauts) underlying a capacity for RNAi appear to have been evolutionarily maintained, a functional divergence among the precise miRNA sequences that trigger gene silencing is evident among species (71). Mechanistic differences in the intercellular transport and systemic distribution of small RNA modifiers of gene expression presumptively represent another logical point of divergence between anatomically distinct organisms. Despite extensive therapeutic interest and success in dsRNA delivery to mammalian cells, it is surprising that systemic mechanisms of small RNA distribution and import in humans remain elusive and largely uncharacterized in the 25+ years since the pioneering discoveries of RNAi and miRNAs were made using *C. elegans* (72-74)

Two mammalian orthologs of *C. elegans* SID-1 encoded within human genome are SIDT1 and SIDT2 (75, 76). Contradictory reports on whether or not these proteins function as dsRNA transporters currently hinder a more definitive classification of their activities. Multiple studies support a role for SIDT1 in dsRNA and miRNA transport in human cells (23, 25, 77). Sidt1 (-/-) knockout mice have also been generated that lack an ability to absorb dietary miRNAs in the gut (78). Sidt2 is highly expressed in several tissues including the brain, intestine, liver, and kidneys of rats (79). Sidt2 (-/-) deficient mice have been associated with impaired glucose tolerance (80), but conversely exhibit reduced tumor burden in lung and gastrointestinal cancer models (25). On the contrary, other reports using HEK293, PANC-1 or *Drosophila* S2 cells, reported that both SIDT1 and SIDT2 lack dsRNA transport activity, and are instead required for the translocation of cholesterol (81, 82). Overexpression of either human SIDT1 or SIDT2, or SID-1 of *C. elegans*, in HEK293 cells revealed that radiolabeled dsRNA uptake displayed a dose-dependent response to cholesterol addition (82). Furthermore, Innate immunity in *C. elegans*, which is a cholesterol auxotroph, requires the function of the CHUP-1 cholesterol transporter, a structural but not functional ortholog of SID-1, as well as human SIDT1 and SIDT2 (83, 84). Likewise, it was demonstrated that SIDT2 functions in the endolysosomal transport of dsRNA from different dsRNA-producing viruses in the initiation of the innate immune response, which is impaired in Sidt2 (-/-) knockout mice (24). Most significantly, a recent report has demonstrated that levels of SIDT2 strongly correlate with those of alpha-synuclein, both of which are increased, in the postmortem brains of patients with PD and Dementia with Lewy Bodies (26). Therefore, the descriptive pathology of human synucleinopathies mirror our functional observations in *C. elegans*; both suggesting that the downregulation of dsRNA transport may be protective to dopaminergic neurons under conditions of neurotoxic stress.

All prior research on SID-3 has focused on its established function in *C. elegans* as a mediator of dsRNA uptake and RNAi silencing in the somatic cells through its regulation of the dsRNA transporter, SID-1 (18). Since *C. elegans* dopaminergic neurons have proven refractory to RNAi in wildtype animals, presumably due to lower levels of SID-1 expression, it was therefore important to clarify whether dsRNA silencing activity was preserved in these cells. Others and we have previously demonstrated that targeted overexpression of *sid-1* in worm neurons, as well as selectively in neuronal subtypes using different promoters, confers enhanced sensitivity to RNAi knockdown by exogenous bacterial feeding of dsRNA (31, 85, 86). Thus, the basic cellular machinery for RNAi is not completely absent from these neurons. It follows that endogenous *sid-1* expression in dopaminergic neurons was confirmed through the recently completed the *C. elegans* neuronal transcriptome (87).

We found that *sid-1(pk3321)* mutant animals displayed significantly less 6-OHDA-induced neurodegeneration when compared to wildtype worms, but this effect was not as strong as was observed with *sid-3(ok973)* mutants (Figure 3C). This is both an interesting and logical result. We demonstrated that *sid-3(ok973)* mutants exhibited neuroprotection from 6-OHDA uptake, as a measure of DAT-1 endocytosis (Figure 1A, B, E). Similarly, we also established that SID-3 modulates RNAi silencing in the dopaminergic neurons, a readout reflective of the endocytic regulation of SID-1 (Figure 3A, B). Thus, the observed neuroprotection in *sid-1(pk3321*) mutants would be expected to be due to a lack of dsRNA import, preventing the silencing of genes involved in pathways required to maintain dopamine homeostasis. In contrast, the enhanced level of protection observed in the *sid-3(ok973)* animals reflects the additive effect that loss of SID-3 function exerts in the co-regulation of DAT-1. We also showed that this regulatory intersection can be modulated through AIM-100 exposures that target and inhibit the activity of SID-3/TNK2. To the best of our knowledge, this is the first time a drug has been shown to impede RNAi silencing, in addition to having neuroprotective activity *in vivo*. Notably, AIM-100 treatments and CRISPR-knockout mutants of *TNK2* have also been shown to prevent viral infection in human cell cultures (20, 88). These combined findings strongly support a conserved functional role for TNK2 as a modulator of dsRNA and illustrate its potential as a druggable target.

Lastly, we sought to provide functional clarity by evaluating the identified PD-related SNPs in *TNK2* (1). Using CRISPR technology, two of these mutations that coded for missense mutations in conserved regions of SID-3 were engineered and the modified animals were assayed for 6-OHDA neurotoxicity and RNAi-targeted knockdown of GFP in dopaminergic neurons. Animals expressing either of the two SNP variants exhibited increased sensitivity to 6-OHDA exposures (Figure 4C). Furthermore, our analyses on the CRISPR-edited animals using either NAB2 or AIM-100 treatment highlighted the pathological importance these variants. The UBA-domain SNP abolished NAB2-protection from 6-OHDA (Figure 4E), whereas animals with the PKD-localized SNP retained neuroprotection with AIM-100 (Figure 4F). This latter result is consistent with what has been shown in human cell cultures where AIM-100 prevented the increased phosphorylation of a TNK2 variant harboring a similar SNP in the same domain (89). Importantly, neither of these SNPs exerted a discernable change in dopaminergic neuronal RNAi silencing (Figure 4D), indicating that SID-3 retained its functional effect on dsRNA import in *C. elegans*. Given that these PD-specific variants appear to encode either a degradation-resistant or hyperactive enzyme, these would not be expected to diminish dsRNA-dependent silencing, as was observed with a *sid-3* knockout mutant (that defines “systemic RNAi deficiency”). Thus, the pathological consequences of this interplay may be significant, and synergistic to effects on dopamine dysregulation, considering that we have shown neuroprotection from 6-OHDA is at least partially an outcome of reduced gene silencing.

Our results highlight the functional significance of both NEDD4 and TNK2, and how their interaction plays a vital role in modulating presynaptic neuroplasticity (Figure 5). Future efforts directed to analysis of the remaining two discovered SNPs in TNK2 are likely to provide further insights (1). Characterization of a mutation in TNK2 (R877H) adjacent to the epidermal growth factor receptor binding domain (EBD; Figure 4A) might be expected to result in a gain-of-function phenotype, since TNK2 phosphorylation was shown to be stimulated by EGFR-ligand binding (89). The other remaining SNP, located in the putative NEDD4-binding motif of TNK2 (Figure 4A; V638M), also warrants attention, given our results with the UBA-localized mutation, and since a similar SNP found in epilepsy patients was reported to block NEDD4-TNK2 interaction (53). Our findings using rat primary neurons (Figure 2), where NAB2 treatment caused a reduction in TNK2 protein levels, in addition to resistance to 6-OHDA neurotoxicity, speak to the translational significance of the interaction between NEDD4 and TNK2 in mammals. Future studies, ideally using mutant *TNK2* patient-derived neurons from iPSCs, would be informative when considering NAB2 was already proven efficacious in conferring neuroprotection to PD-patient derived dopaminergic neurons in culture (8).

The optimal function and survival of dopaminergic neurons is a consequence of a variety of intrinsic factors (genetics, proteostasis, aging, oxidative stress, mitochondrial/lysosomal dysfunction), all of which combine to modulate brain activity. Given the scale and diversity of dopamine-associated disorders, one would not be incorrect to assume these factors seemingly act near a precipice of dysfunction – yet are poised for response to epigenetic influences that raise or lower this threshold. That living organisms have evolved integrated cellular mechanisms to provide for a capacity to respond, adapt, and buffer neurons to and from external stressors is as fragile as it is elegant. No finer example of the delicate balance between resilience or susceptibility, and truly life or death for some, can be found than what exists in the intricate management of dopaminergic neuron health.

## Materials and Methods

### *C. elegans* 6-OHDA Neurodegeneration Assays

Animals were scored for 6-OHDA-induced neurodegeneration as previously described (39, 40). Briefly, worm strains were synchronized at hatching and raised on agar plates of Nematode Growth Media (NGM) seeded with *E. coli* strain OP-50 at 20°C. L4-stage larvae were collected from plates by resuspension using ddH_2_O and transferred to 10mL glass conical tubes, centrifuged, and then washed thrice with ddH_2_O to remove the bacteria. Animals were subsequently transferred to 1.5mL Lo-Bind tube and treated with 6-hydroxydopamine (6-OHDA; Tocris, Bristol, UK) containing 2mM ascorbic acid for one-hour, while rotating at 20°C. After the one-hour incubation with 6-OHDA, M9 was added to oxidize the 6-OHDA and terminate the experiment. Worms were immediately washed thrice with ddH_2_O or until the supernatant was no longer pink (indicative of oxidation). Washed worms were placed onto freshly seeded plates and scored for neurodegeneration twenty-four hours later.

The six anterior dopaminergic neurons of *C. elegans* were scored in each animal as previously described (38, 39). Briefly, worms were scored under a fluorescent microscope using the 60X objective. Worms were scored as “normal” if no degenerative phenotypes (dendritic blebbing, swollen soma, broken projections, shrunken soma, or a missing neuron) in the six anterior [four CEPs (cephalic) and two ADEs (anterior deirid)] dopaminergic neurons. In total, three trials consistent of 30 animals per replicate, totaling to 90 animals per strain/condition.

### AIM-100 & NAB2 Drug Treatments in Neurodegenerative Assays

Stocks of AIM-100 and NAB2 (Tocris; Bristol, UK) were dissolved in 200 proof ethanol and kept at 4°C. Synchronized *C. elegans* were treated daily with either drug. For drug exposure, worms were resuspended from OP-50 plates with ddH_2_O, transferred to 10mL glass conical tube, and washed thrice with 10mL of ddH_2_O to remove bacteria. Worms were then transferred to a fresh 1.5mL Lo-Bind tube and suspended in 100µM concentration of AIM-100, NAB2, or 0.01% ethanol diluted in 0.5X M9. 1.5mL Lo-Bind tubes of worms and drugs were kept rotating at 20°C for one-hour. After exposure, drugs were washed away thrice with ddH_2_O, and worms were placed onto fresh OP-50 plates, then maintained at 20°C. On days of 6-OHDA treatments, worms were given the drug treatment (either AIM-100 or NAB2) as described, transferred to fresh 1.5mL Lo-Bind tube after the drug has been washed away, then immediately exposed to 6-OHDA. Twenty-four hours later the animals were scored for neurodegeneration.

### *C. elegans* RNAi

The GFP RNAi construct was plasmid L4417 was a gift from Andrew Fire (Addgene plasmid #1649). Bacteria were isolated and grown overnight at 37°C shaking in Luria broth (LB) containing 100µg/ml ampicillin. NGM plates containing 50µg/ml ampicillin and 1mM IPTG were seeded with 300µl of RNAi bacterial culture and allowed to dry and grown overnight at 20°C under foil. Animals were raised on RNAi bacteria for two generations and scored for dopaminergic GFP silencing at day 5 post-hatching in the second generation. The six anterior dopaminergic neurons were scored under a fluorescent microscope using a 60X objective. If all six anterior neurons were visible, then the animal was scored as “normal” while if any of the neurons were not visible (i.e., non-fluorescent) then the animal was scored as abnormal.

### Swimming-Induced Paralysis (SWIP)

SWIP analysis was performed as previously described (43). Briefly, 10-15 worms, synchronized to the L4 stage, were placed into well containing 200µL of water and allowed to thrash for 10 minutes. After 10 min., the wells were observed by eye under a dissecting scope and the number of animals still moving were recorded. Ten wells were scored for each strain. Significance was determined by one-way ANOVA with Dunnett’s post hoc test.

### *C. elegans* Strains

Worms were grown on Petri plates containing standard Nematode Growth Medium (NGM) that were seeded with *E. coli* strain OP-50 bacterial lawns. All strains were maintained at 20°C under standard laboratory conditions (90). Strains used in this study include: N2 (Bristol; “wildtype”), NL3321 (*sid-1(pk3321))*, VC787 (*sid-3(ok973))*, RM2702 *(dat-1(ok157))*, GR1373 (*eri-1(mg366))*, VC889 *(wwp-1(gk372))*, PHX898 *(sid-3(syb898))*, PHX912 *(sid-3(syb912)*, and BY250 (*vtIs7*[P_*dat-1*_::GFP]; kindly provided by Randy Blakely). Crosses performed to generate strains used in this study are in Table 1. Briefly, BY250 was crossed separately into NL3321, VC787, RM2702, GR1373, VC889, PHX898 and PHX912 to create UA423, UA424, UA407, UA425, UA426, UA427 and UA428, respectively (Table 1). The resulting cross of UA425 was further crossed separately into both PHX898 and PHX912 to create UA429 and UA430, respectively (Table 1). Genome editing of *C. elegans* (wildtype, N2) was performed to generation the *sid-3(syb898) and sid-3(syb912)* mutant alleles, strains PHX898 and PHX912, respectively, using the CRISPR technique by SunyBiotech (Fujian, China). The specific sgRNAs used in creating PHX898 were: 5’-CCCAAGTACTGCTCGGAGCG-3’, and 5’-GAATTGTTGGAAATTCAATC-3’; and for PHX912: 5’-CAGTTAAATGGATATCCAAA -3’; 5’-AACGGGTACCCTCAGTTAAA<u>TGG-</u>3’, and synonymous mutations were introduced into the PAM sites. PHX898 contains a single nucleotide polymorphism in the *sid-3* gene encoding a I341A change in the SID-3 protein, located in the protein kinase domain. PHX912 contains a single nucleotide polymorphism encoding a A1016V change in SID-3 located in the ubiquitin-binding association region of this protein.

**Table 1.** Completed Crosses for Strains Used

| **Strain** | **Genotype** | **Strains Crossed** |
| --- | --- | --- |
| UA423 | *sid-1(pk3321); vtIs7[P<sub>dat-1</sub>::GFP]* | NL3321 x BY250 |
| UA424 | *sid-3(ok973); vtIs7[P<sub>dat-1</sub>::GFP]* | VC787 x BY250 |
| UA407 | *dat-1(ok157); vtIs7[P<sub>dat-1</sub>::GFP]* | RM2702 x BY250 |
| UA425 | *eri-1(mg366); vtIs7[P<sub>dat-1</sub>::GFP]* | GR1373 x BY250 |
| UA426 | *wwp-1(gk372); vtIs7[P<sub>dat-1</sub>::GFP]* | VC889 x BY250 |
| UA427 | *sid-3(syb898); vtIs7[P<sub>dat-1</sub>::GFP]* | PHX898 x BY250 |
| UA428 | *sid-3(syb912); vtIs7[P<sub>dat-1</sub>::GFP]* | PHX912 x BY250 |
| UA429 | *sid-3(syb898); eri-1(mg366); vtIs7[P<sub>dat-1</sub>::GFP]* | PHX898 x UA425 |
| UA430 | *sid-3(syb912); eri-1(mg366); vtIs7[P<sub>dat-1</sub>::GFP]* | PHX912 x UA425 |

### Culture of primary neurons

Primary rat cortical and dopaminergic neurons were prepared from rat feti (Sprague-Dawley, day 18 of gestation; Envigo, Indianapolis, IN) as described previously (91). Briefly, neurons (0.5 × 10^6^ cells/ 35mm plate) were grown in neurobasal medium supplemented with B-27, glutamine, and antibiotics (Thermo Fisher Scientific, Waltham, MA, USA) for 12 days in vitro. NAB2 treatment: A solution of NAB2 (Tocirs, Bristol, UK) was prepared in filter-sterilized 99.9% DMSO (Sigma-Aldrich, St. Louis, MO, USA) and added to the cell culture medium (final concentration: 2 mM) for 24 hours, at which point they were analyzed. The vehicle control group was treated with isovolumetric sterile DMSO. 6-OHDA treatment: 6-OHDA (final concentration 20μM) was prepared in sterile water and added to the cell culture medium. The stock solution of 6-OHDA was prepared in ascorbic acid (final concentration 4 μM). The vehicle control group for 6-OHDA was treated with isovolumic sterile water. Cells were treated for 24 hours and then immediately analyzed. All protocols were approved by the Institutional Animal Care Committee (IACUC) of The University of Alabama, Tuscaloosa, AL.

### Immunoblots

After treatment with or without NAB2 for 24h, cortical neurons were scraped and lysed in 1X cell signaling buffer (Cell Signaling Technology, Danvers, MA, USA) and protein concentration was determined using the BCA protein assay (Thermo Fisher Scientific, Waltham, MA, USA). Samples (50µg of protein/lane) were separated on a 4-12% SDS-polyacrylamide gel (Bio-Rad, Hercules, CA, USA) and probed with anti-ACK antibody (1:100, Santa Cruz, Dallas, TX, USA) and anti-β-actin (1:1000, Sigma-Aldrich, St. Louis, MO, USA). Scanned images were analyzed using ImageJ software (National Institutes of Health, Bethesda, MD, USA).

### Immunocytochemistry

Primary dopaminergic neurons fixed in 10% buffered formalin were blocked in 10% goat serum for 1 h, then incubated with anti-tyrosine hydroxylase (TH) antibody (1:100, Pelfreez-Bio, Rogers, AR, USA) overnight at 4 °C. Cells were washed and incubated with Alexa-568 antibody (1:200 dilution; Invitrogen, Molecular Probes, Carlsbad, CA, USA) for 1 h at room temperature. Images were taken with a Zeiss Axiovert A1 microscope. The number of TH-positive live cells and TH-positive dead cells was counted using AxioVision 4.9. The viable dopaminergic neurons percentage was calculated using the following equation: Viable TH-positive cells (%) = the number of TH-positive and DAPI-positive live cell / (TH-positive and DAPI-positive live cells + TH-positive and DAPI-positive dead cells) x 100.

## Acknowledgments and Funding Sources

We wish to acknowledge all members of The Caldwell Lab for their teamwork and collegiality. Special thanks are in order for the technical assistance of Adam Holzhauer and Brian Smithers, as well as the expert advice and support of Laura Berkowitz. We are especially grateful to Anthony Gaeta, Edward Griffin, Jennie Thies, and Xiaohui Yan for their scientific insights. We gratefully acknowledge Randy Blakely and Andrew Fire for strains and vectors; we wish to also thank WormBase. This research was funded by a grant to Guy A. Caldwell from the U.S. Department of Health & Human Services, National Institute of Neurological Disorders and Stroke (R15NS104857). Some strains were provided by the CGC, which is funded by NIH Office of Research Infrastructure Programs (P40 OD010440); other support came from The University of Alabama College of Arts & Sciences, The Hill Crest Foundation, and Parkinson’s Disease Support Group of Huntsville, Alabama.

